# Habitat specialisation and dispersal capacity drive rapid carabid beetle responses to urban forest fragmentation

**DOI:** 10.1101/2025.10.17.683013

**Authors:** Basile Finand, D. Johan Kotze

## Abstract

The effects of habitat fragmentation on insects are well documented, yet most studies overlook extinction debt. We investigated carabid beetles in 25 remnant urban forests in Helsinki, Finland, spanning unfragmented, recently fragmented, and historically fragmented sites. Across 3162 individuals and 34 species, we analysed species richness, community composition, and traits including dispersal capacity, habitat specialisation, and body size both at the community and population levels. We found no extinction debt: species richness remained stable, but community composition shifted rapidly. Forest specialists declined non-linearly within three decades post-fragmentation before partially recovering, whereas open-habitat species showed the opposite pattern. Dispersal-limited species lost richness without compositional change, while highly-dispersive species maintained richness but altered community composition. Individual size, mass, and dispersal traits showed no consistent patterns. Our study demonstrates rapid, trait-mediated responses to fragmentation in short-lived beetles and highlights the importance of considering specialisation and dispersal in urban conservation planning.

## Introduction

Human activities significantly affect ecosystems, with 75% of terrestrial environments considered severely degraded (IPBES 2018; Venter *et al*. 2016). A main driver is habitat fragmentation, defined as the division of large, continuous habitat into smaller, more isolated patches (Fahrig 2003, 2017). Fragmentation can result from urbanisation, agriculture, or transport infrastructure and has significant, multifaceted consequences for biodiversity. It reduces biodiversity by 13 to 75%, depending on the location and ecosystem considered (Haddad *et al*. 2015), and disrupts ecosystem functioning, affecting processes such as nutrient cycling and food webs. Moreover, it can select for species traits, including behaviour, reproductive strategies, body size and dispersal capacities (Bernardino *et al*. 2024; Cheptou *et al*. 2008; Finand *et al*. 2023; Kurki *et al*. 2000; Mahan & Yahner 1999; Warzecha *et al*. 2016). By altering dispersal abilities, fragmentation modifies gene flow and, in the longer term, can influence evolution (Young *et al*. 1996).

At the community level, many studies emphasise the negative consequences of habitat fragmentation and loss on taxa, particularly in terms of species richness and community composition. For example, there are fewer ant species in small urban parks compared to larger ones in Spain (Carpintero & Reyes-López 2014), or fewer anuran species in fragmented forests in the Brazilian Cerrado (Ramalho *et al*. 2022). Similar patterns have been reported for birds (Herkert 1994), butterflies (Soga & Koike 2012), or mammals (Crooks 2002). Active debates persist regarding the actual effect of fragmentation *per se* compared to the effect of habitat reduction or loss (Fahrig 2017; Riva *et al*. 2024). Some studies suggest that, for the same amount of habitat, fragmentation could even have a positive effect (Perrin *et al*. 2025), although this is not always supported (Gonçalves-Souza *et al*. 2025). Here, we consider habitat fragmentation as the overall process, integrating both increased isolation and habitat loss; we do not focus on fragmentation *per se*.

While most studies focus on current habitat configuration, the importance of time in ecology, and especially in urban ecology, is receiving increasing attention (Chen *et al*. 2023; Ossola *et al*. 2021). A time lag can occur before the effects of habitat fragmentation on biodiversity become apparent, a phenomenon known as the extinction debt (Kuussaari *et al*. 2009; Tilman *et al*. 1994). The length of this time lag can depend on several factors. Theoretical work has shown that the lag can be longer when populations are close to their extinction threshold (Hanski & Ovaskainen 2002). Species traits are also important: short-lived, poor-dispersing, and specialist species are likely to exhibit a shorter time lag following habitat changes (Chen *et al*. 2023). The study of this time lag is especially important in the context of urbanisation, where changes happen quickly, making it difficult for species to adapt. We aim to understand the impact of time on communities of insects in the context of urbanisation.

Some studies have demonstrated this extinction depth, where past habitat configuration explains current communities better than the present landscape. Many of them have focused on plants (Cousins 2009; Helm *et al*. 2005; Lindborg & Eriksson 2004; Löffler *et al*. 2020), showing that it depends on traits such as longevity (Krauss *et al*. 2010) or habitat type (du Toit *et al*. 2016). This time lag can be long, sometimes exceeding 100 years (Hahs *et al*. 2009; Vellend *et al*. 2006). A few studies have examined this phenomenon in other taxa such as birds (Almeida-Gomes *et al*. 2025), mammals (Ancillotto *et al*. 2025), and pollinators (Cusser *et al*. 2015; Löffler *et al*. 2020; Soga & Koike 2013). However, little attention has been given to insect groups other than pollinators, especially less mobile ones. We aim to fill this gap by studying the response of carabid beetles to forest fragmentation time lags in the urban milieu.

Carabid beetles are an appropriate group to study habitat fragmentation and the importance of time. It is a species-rich group occurring in many different habitat types (Lövei & Sunderland 1996). They harbour variation in life history traits of interest, such as body size, dispersal strategies (presence/absence of wings), habitat preference with different degrees of specialism, or food preferences (Lindroth 1985, 1986). Moreover, studies have demonstrated the impact of current habitat fragmentation on their communities (Fujita *et al*. 2008; Koji *et al*. 2024; Niemelä 2001). At the population level, habitat fragmentation and urbanisation also affect morphological traits (Keinath *et al*. 2023; Kotze *et al*. 2024; Papp *et al*. 2020; Weller & Ganzhorn 2004). Studying carabid beetles allows us to fill the gap on the importance of extinction debt on insects, and the particular importance of their different life history traits.

Our study aims to understand how the age of habitat fragmentation impacts carabid beetle communities and populations by sampling remnant urban forests in the Helsinki metropolitan area in Finland. We expect a decrease in species richness with an increase in the age of fragmentation. Moreover, we predict that community composition will differ depending on age. Because carabid beetles are short-lived (a few years), we hypothesise that the extinction debt will be short. However, we propose that wingless and specialist species will be the most impacted, with a quicker decrease in richness and change in community composition. At the population level, we project a decrease in the size and mass of the beetles within the fragmented sites due to the urban heat island effect and urban disturbances, this decrease being more pertinent in the historically fragmented sites (Craig Stillwell & Fox 2009; Sheridan & Bickford 2011; Tseng *et al*. 2018).

## Material and methods

### Study area and sites

The study was conducted in the Helsinki metropolitan area, southern Finland (60°10′15″N, 24°56′15″E), during the summer of 2023. We selected 25 sites in three categories (treatments) of fragmentation age: five control forests with no significant fragmentation, ten recently fragmented forests, where fragmentation occurred within the last 35 years, and ten historically fragmented forests, fragmented more than 35 years ago (Fig. 1, Table S1). Site abbreviations are explained in Table S1: if the site code starts with a C, it represents a control site, if it starts with an H = historically fragmented site, R = recently fragmented site. The forests were selected using QGIS and historical aerial photographs of the Helsinki region (City of Helsinki, https://kartta.hel.fi/). Sites were spruce-dominated, ≥1 ha, and ≥1 km from the Baltic Sea. We digitised all the remained forests in Helsinki for three time points: 2023 (year of sampling), 1988 (marking the ∼35-year threshold), and 1932 (earliest available aerial photographs). For each forest present in 2023, we determined when fragmentation occurred, whether it happened before or after 1988. We considered a forest to be fragmented when it had become disconnected from a larger forest and consequently reduced in size. In this study, ‘habitat fragmentation’ thus refers to both a reduction in size and isolation from other forested areas. In addition to the categorical age of fragmentation (control, recently, historically), we calculated a rough numerical age of fragmentation. Thanks to the availability of aerial images at roughly ten-year intervals, we recorded the numerical fragmentation age as the first year in which the forest appeared disconnected. As a result, the numerical age of fragmentation in this study is accurate to within approximately zero to nine years. The five control sites consist of large forests around the city that have largely remained unfragmented. Three of these control forests (site codes CSIP, CTUU, CNUU) are extensive forests or national parks located outside the city, while two (CESP, CPAL) are large forests situated within the urban environment (Fig. 1).

**Figure 1:**
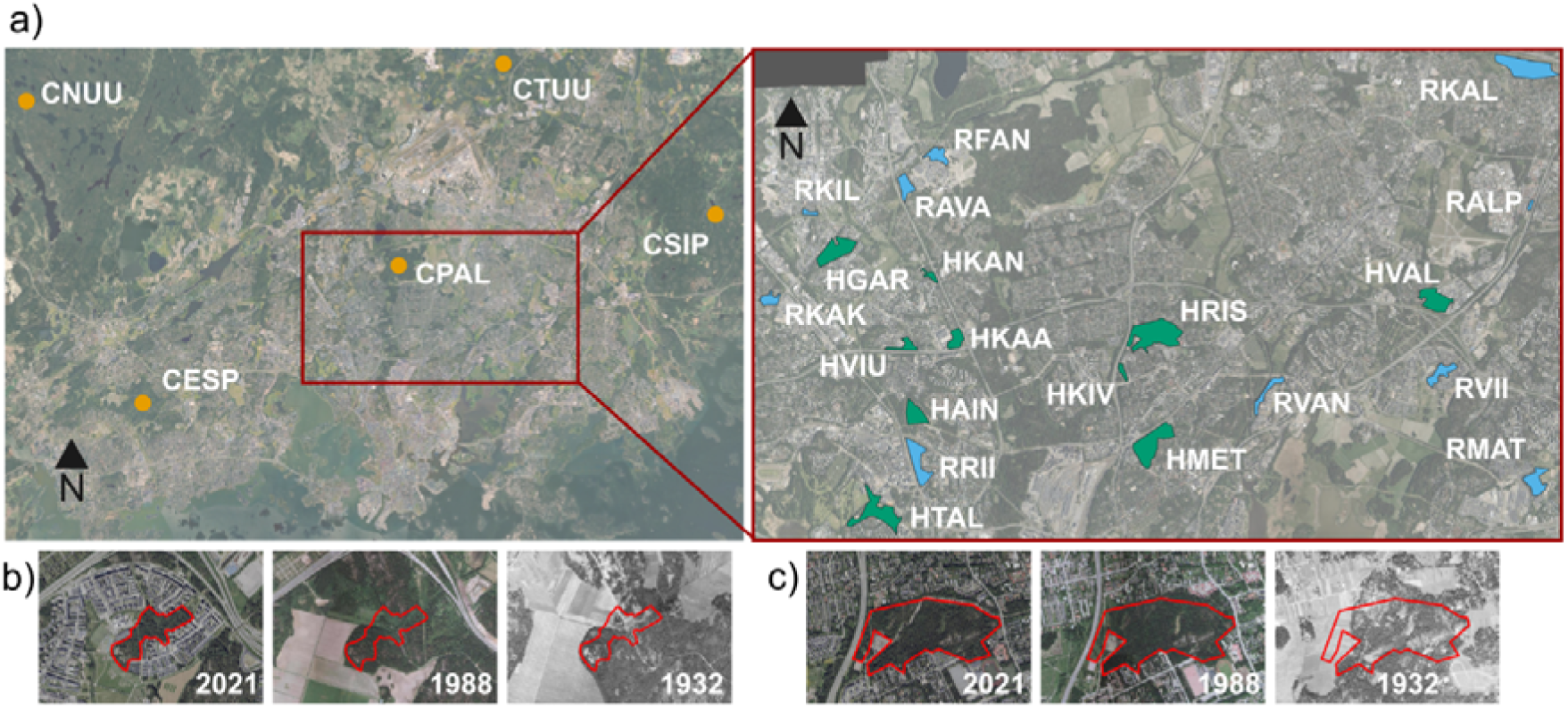
a) Map of the control sites (left panel) and urban sites (right panel). Blue sites are the recently fragmented sites, and green sites the historically fragmented sites. b) Example of a recently fragmented site (RVII) when fragmentation occurred between 1988 and 2021. c) Example of a historically fragmented site (HRIS) when fragmentation occurred between 1932 and 1988. Source of the aerial pictures: Helsinki region provided by the City of Helsinki (https://kartta.hel.fi/).

### Carabid beetle sampling and identification

We sampled ground beetles between 5 June and 11 October 2023 using pitfall traps. At each site, we installed ten pitfall traps arranged in two lines 5 m apart, with traps per line also 5 m apart. Each trap consisted of a plastic cup (65 mm in diameter), covered by a plastic roof to prevent water and debris from entering. The cups were half-filled with a 50% propylene glycol/water solution and embedded in the soil so that the rim was level with the ground surface. Traps were checked approximately every three weeks, resulting in six visits in total. During each visit, we emptied the contents of the traps, preserved the collected specimens in alcohol until laboratory identification, and refilled the traps with propylene glycol.

In the laboratory, we sorted the traps’ contents and retained only the carabid beetles. The catch of all traps per site and across all visits was pooled for analysis. Individuals were identified to species level using Lindroth (1985, 1986). We recorded wing morphology (short or long) and measured the elytra length under a binocular microscope. Each individual was weighed after air-drying for one week.

Species-level traits, such as habitat, humidity, and feeding preference, were compiled from Lindroth (1985, 1986). For wing morphology and body size, we used measurements from collected specimens to calculate mean body size and to determine the wing morphology of each species.

### Environmental measurements

At each site, we measured environmental characteristics (Table S1). We took five canopy photos (at traps 1, 3, 5, 7, and 9) by placing a camera on a 10 cm high plastic box. Images were binarised in ImageJ to count the number of white pixels, representing the sky, as an indicator of canopy openness. We calculated the average canopy openness for each site. We measured litter depth at two locations around each trap, resulting in 20 measurements per site. This was done using a nail inserted into the litter until resistance indicated the underlying soil. We then averaged these measurements per site. Using a 1 × 1 m quadrat, we visually estimated the percentage cover of the field and ground layer at five locations (traps 1, 3, 5, 7, and 9) and calculated the mean value per site. In two 10 × 10 m squares centred on traps 3 and 7, we measured the total area of paths within each square, recording the sum of all path areas (in m²). Within these same squares, we counted all trees with a diameter at breast height (DBH) greater than 10 cm. We also measured the total length and mid-point diameter of all logs on the ground with a diameter greater than 5 cm. Log volume was calculated using the formula for a cylinder: π × length × (diameter/2)². We similarly recorded the height and average diameter of stumps and calculated their volume using the same method. We then summed the volumes of logs and stumps to obtain the total volume of deadwood per site. At each site, we also installed two temperature loggers (at traps 3 and 7), placed 3-5 cm below the soil surface, which recorded temperatures every hour throughout the sampling period and the subsequent winter until 5 May 2024. We calculated the mean temperature per site from these two loggers. For sites where data were lost from one logger (CESP, CNUU, CTUU, HAIN, HKAN, HVAL, RFAN, RKAK, RVAN, and RVII), we used the remaining logger for the affected period. In HKIV, both loggers failed during winter: we estimated the mean annual temperature by modelling the relationship between mean annual and summer temperatures from the other sites with a linear model (correlation: 73%) and applying the parameters of the model to the summer temperature of HKIV to estimate mean annual temperature. Minimum and maximum temperatures were highly correlated with mean temperature. So, we decided to only use mean temperature.

Finally, we collected soil samples at each trap, storing them in one plastic bag per site. We measured soil pH, organic matter content, and moisture in the laboratory. To prepare the samples, we first sieved the contents of each bag using a 2 mm sieve to homogenise the soil and remove roots, stones, and other debris. The sieve was thoroughly washed between samples to avoid cross-contamination. For soil moisture and organic matter analysis, we placed 1 dl of sieved soil from each sample into an aluminium cup. We recorded the mass of the empty cup and then the mass of the cup with the soil before placing it in an oven at 100 °C overnight. The following morning, we weighed the dried samples and calculated the percentage of soil moisture. For organic matter analysis, we first weighed ceramic crucibles. The dried soil samples were ground using a mortar and pestle, and approximately two teaspoons of soil per site were placed into each crucible (one crucible per site). We recorded the mass of the filled crucibles before placing them in a muffle furnace (Umega SNOL 30/1100 with Omron E5CK-T controller) at 550 °C for 4 h. After cooling the crucibles in a desiccator, we weighed them again to calculate percentage organic matter. Finally, to measure soil pH (pHmeter WTW inoLab pH level1), we transferred 15 ml of sieved soil into 50 ml plastic tubes and added 30 ml of ultrapure water. The mixture was vortexed for 10 s and allowed to settle for 1 h. Afterwards, the samples were centrifuged for 5 min, and the pH of each supernatant was measured.

### Statistical analysis

Data analyses were carried out using R version 4.2.2 (R Core Team 2022). Throughout all analyses, a p-value < 0.05 was considered to indicate a significant effect, and a p-value < 0.1 an indicative or slight (marginal) effect.

To verify that our sites were of the same type, i.e., differed only in fragmentation age but were otherwise comparable for the other environmental variables measured, we performed a principal component analysis (PCA) and tested for statistical differences using a permutational multivariate analysis of variance (PERMANOVA) with distance matrices, implemented via the *adonis2* function in the vegan package (Oksanen *et al*. 2022).

To select uncorrelated environmental variables for inclusion in subsequent models, we conducted pairwise correlation analyses. For pairs of numerical variables, we used Spearman’s correlation tests; for pairs involving one numerical and one categorical variable, we applied Kruskal–Wallis tests; and for pairs of categorical variables, we used Fisher’s exact tests. We then retained the variables that were not correlated with treatment (control, historically fragmented and recently fragmented sites) or with each other.

To assess the influence of environmental variables on total species richness, we fitted linear models. We also calculated the cumulative number of species per treatment using the *iNEXT* function (Hsieh *et al*. 2024). To investigate the role of traits in mediating the effect of fragmentation age on species diversity, we divided the data into five subsets: short-winged species, long-winged species, forest specialists, open-habitat specialists, and generalists. These traits were hypothesised to be highly responsive to fragmentation history (Chen *et al*. 2023). We then fitted either linear or polynomial models, depending on the structure of the data.

To compare community composition among treatments and to assess the effects of environmental variables, we performed a distance-based redundancy analysis (db-RDA) using the *capscale* function in the vegan package (Oksanen *et al*. 2022). When treatment effects were significant, we conducted pairwise comparisons using the *multiconstrained* function from the BiodiversityR package (Kindt & Coe 2005).

To examine the relationships between beetle traits and environmental variables, we carried out a fourth-corner analysis using the ade4 package (Thioulouse *et al*. 2018).

Finally, at the population level, we tested the effect of treatment on the size, relative mass (corrected by size), and relative wing length (corrected by size) of individuals belonging to the most abundant species: *Amara brunnea*, *Calathus micropterus*, *Carabus hortensis*, *Carabus nemoralis*, *Cychrus caraboides*, *Leistus ferrugineus*, *Notiophilus biguttatus*, *Pterostichus melanarius*, *P. niger*, and *P. oblongopunctatus*. We used linear models (LM) when assumptions were met, or Kruskal–Wallis tests when they were not.

For all models (LM, GLM, db-RDA), we followed the same procedure to identify the best-fitting model. First, we fitted a model including all uncorrelated environmental variables. Next, we checked model assumptions, such as normality and homoscedasticity, using the DHARMa package (Hartig 2022). We then calculated the Akaike Information Criterion (AIC) of the model. We sequentially removed non-significant variables, one at a time, until we obtained the lowest AIC. This final model was then used to test significance of the remaining variables. For count data, such as species richness, we also used generalised linear models (GLMs) with a Poisson distribution. However, these models consistently yielded higher AIC values than the corresponding linear models, and no issues of normality or homoscedasticity were detected in the linear models.

## Results

We collected a total of 3162 carabid beetle individuals belonging to 34 species (Table S2). The most collected species were *Amara brunnea* (1094 individuals), *Calathus micropterus* (469), *Trechus secalis* (326), *Pterostichus niger* (286), *Carabus nemoralis* (180), and *Pterostichus oblongopunctatus* (176).

The PCA and permutational analysis of variance (Adonis; F = 0.718, p = 0.587) confirmed that the sites were highly similar overall, indicating that the forests primarily differed in fragmentation age (Fig. S1). A few environmental variables, such as canopy openness and litter depth, showed some correlation with treatment (canopy openness: Kruskal-Wallis, χ² = 6.142, p = 0.046; litter depth: Kruskal-Wallis, χ² = 6.142, p = 0.046).

Based on the correlation analyses (Fig. S1), the final set of environmental variables retained for inclusion into models was treatment, percentage organic matter, % field layer cover, % ground layer cover, path surface area, deadwood quantity, and mean temperature.

### Overall species

Total species richness did not differ significantly between treatments (LM, F = 0.974, p = 0.394; Fig. 2). Mean temperature positively influenced species richness (LM, F = 9.329, p = 0.006, Fig. 2), while % field layer cover showed a marginally negative effect (LM, F = 3.694, p = 0.069). Extrapolated species numbers revealed that the historically fragmentation sites supported more species than the recent sites, which in turn had more than the control sites (Fig. 2).

**Figure 2:**
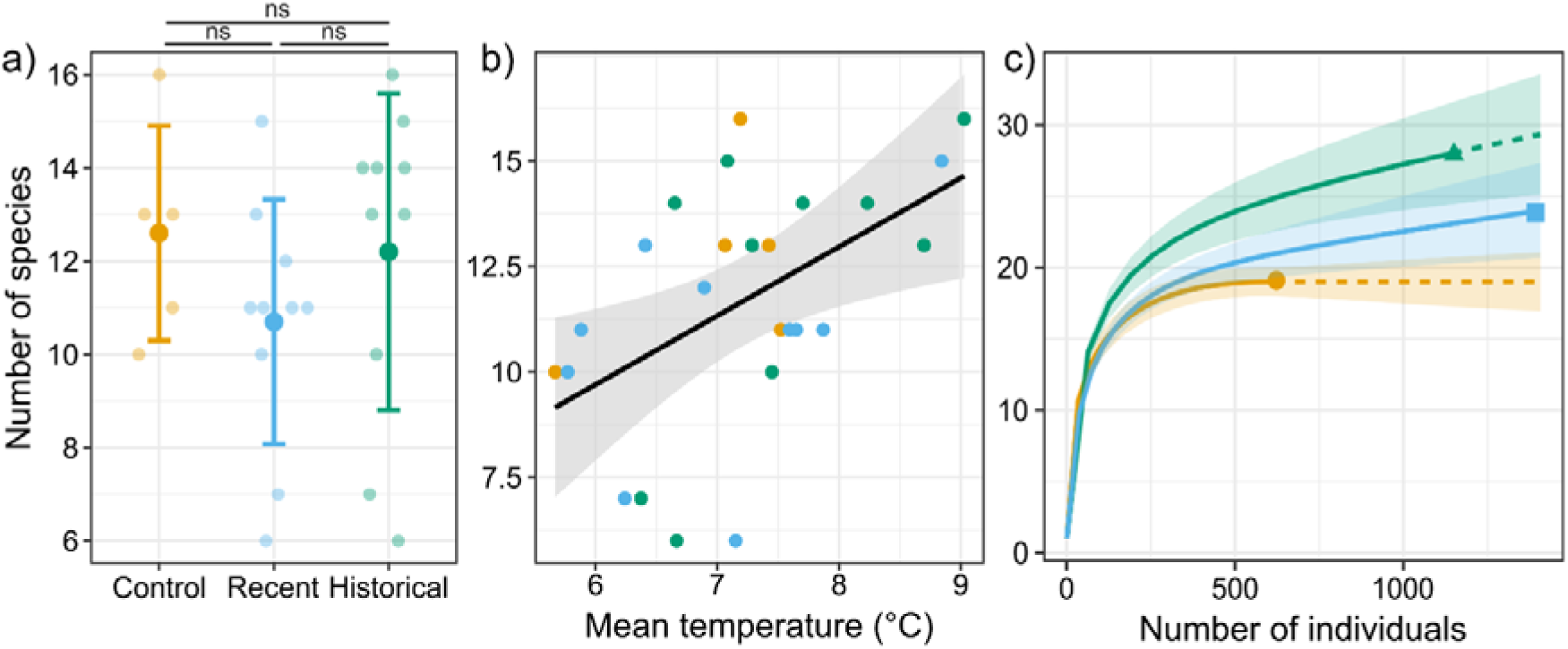
Number of species (± SD) as a function of: a) treatment and b) mean temperature. Orange points represent control sites, blue points represent recently fragmented sites, and green points represent historically fragmented sites. c) Species accumulation curves (solid lines) and extrapolations (dashed lines) by treatment. Orange lines correspond to control sites, blue lines to recently fragmented sites, and green lines to historically fragmented sites.

Only treatment had a significant effect on the community composition of all species (db-RDA, F = 2.469, p = 0.005, constrained variance explained = 0.183, Fig. 3). Differences were found between control and historical sites, and control and recent sites (pairwise comparisons: control-historical, F = 2.574, p = 0.008; control-recent, F = 4.05, p = 0.003), while no significant difference occurred between recent and historical sites. Note that the two urban control sites (CPAL, CESP) are closer to the other fragmented sites than the three more rural control sites. Also, the historical sites displayed greater variation than the recent sites.

**Figure 3:**
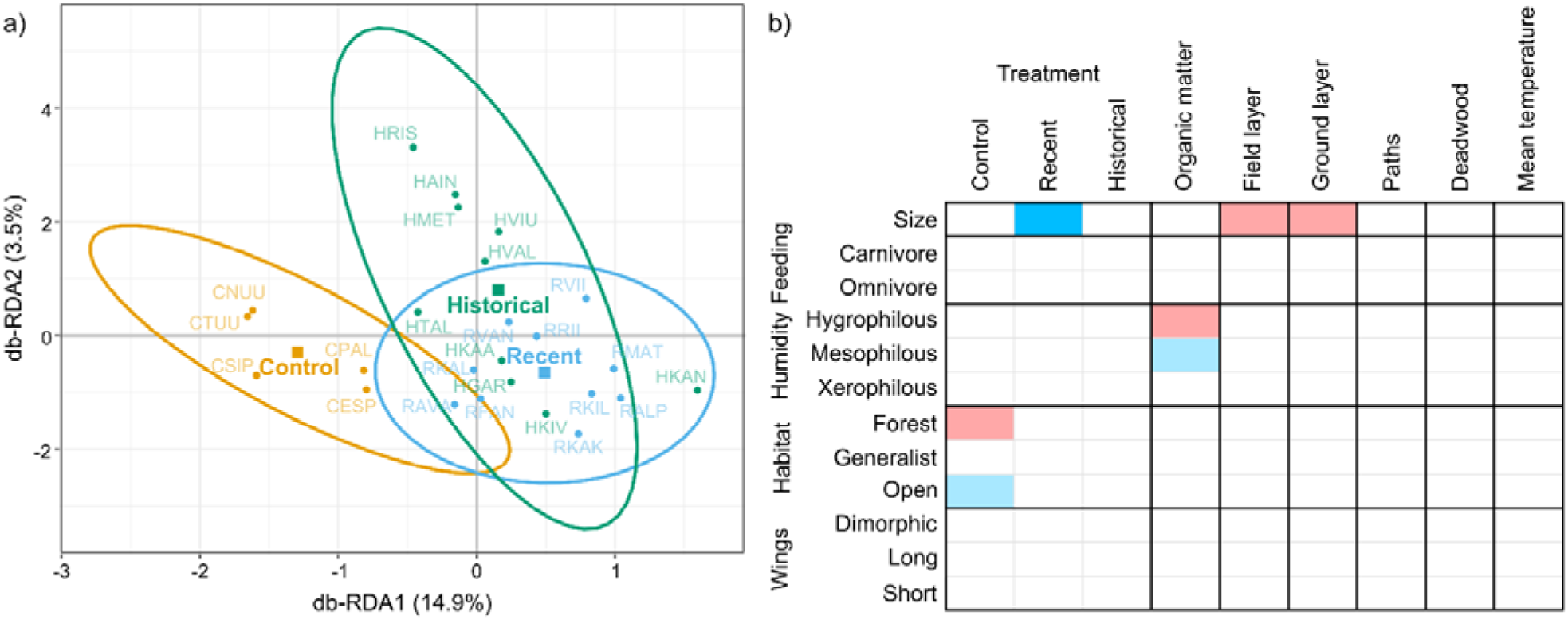
a) Distance-based Redundancy Analysis (db-RDA) showing the effect of treatment. Percentages on the axes indicate the proportion of variance explained by the constrained axes. Ellipses represent 95% confidence intervals. b) Fourth-corner analysis linking species traits with environmental variables. Colours indicate the strength and direction of associations: red denotes positive correlations and blue negative correlations. Light shades represent indicative associations (p < 0.1), while dark blue highlights significant negative correlations (p < 0.05).

In terms of traits at the community level, species were significantly smaller in recently fragmented sites (fourth corner test, p = 0.021, Fig. 3). There was an indicative association between control sites and a higher proportion of forest habitat species (p = 0.063) and fewer open habitat species (p = 0.094, Fig. 3). Larger species were also linked to increased percentages of field (p = 0.066) and ground layer covers (p = 0.063, Fig. 3). Additionally, higher organic matter percentages corresponded to an indicative increased proportion of hygrophilous species (p = 0.09) and a decreased proportion of mesophile species (p = 0.09).

### Habitat specialist species

For forest specialist species, both treatment (LM, F = 3.089, p = 0.069, Fig. 4) and path surface area (LM, F = 3.39, p = 0.081) had indicative positive effects, particularly between control and historical sites. Modelling fragmentation age as a continuous variable revealed a better fit with a polynomial model (AIC linear = 110, AIC polynomial = 106), showing a non-linear pattern: species richness declined until around 30 years post-fragmentation before increasing again (Polynomial, F = 3.344, p = 0.054, R² = 0.163; Fig. 4).

**Figure 4:**
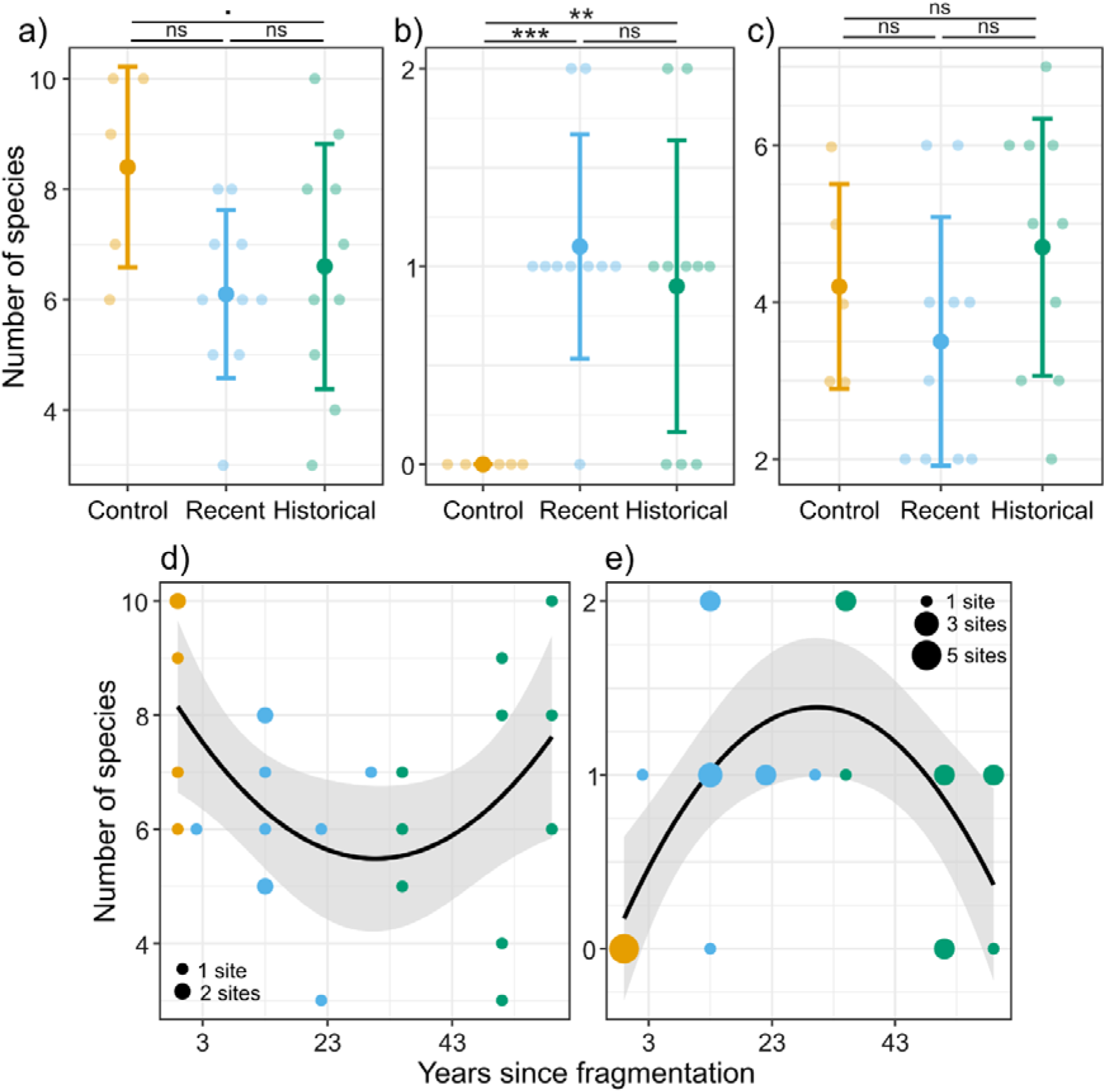
Number of species (± SD) per treatment for a) forest specialist species, b) open-habitat species, and c) generalist species. Number of species related to the number of years since fragmentation for d) forest specialist species, and e) open habitat species. Orange, blue, and green points are, respectively, the control, recent, and historical sites. The size of the points represents the number of sites with this value.

Open habitat specialists were significantly affected by treatment (LM, F = 11.709, p < 0.001, Fig. 4), with more species in the fragmented urban forests than the controls, and positively by deadwood quantity (LM, F = 6.446, p = 0.02). The polynomial model again provided a better fit for age of fragmentation effects on richness (AIC linear = 58, AIC polynomial = 47), showing an increase up to 30 years after fragmentation followed by a decline (Polynomial, F = 7.306, p = 0.004, R² = 0.345; Fig. 4).

Generalist species richness was unaffected by treatment (LM, F = 11.776, p = 0.003, Fig. 4) but positively influenced by mean temperature. Organic matter percentage (LM, F = 3.463, p = 0.078) and % field layer cover (LM, F = 4.3, p = 0.051) had marginal positive and negative effects, respectively. Unlike specialists, a linear model better explained the relationship with fragmentation age (AIC linear = 98, AIC polynomial = 99), though this effect was not significant (LM, F = 0.951, p = 0.34).

### Dispersal capacities

Examining dispersal capacity, treatment had no effect on long-winged species richness (LM, F = 1.177, p = 0.327; Fig. 5) but influenced community composition, with significant differences between control and fragmented sites (db-RDA, F = 3.309, p = 0.002; pairwise: control-recent p = 0.001, control-historical p = 0.004; Fig. 5). Conversely, short-winged species richness was reduced in fragmented sites regardless of fragmentation age (LM, F = 5.477, p = 0.012, R² = 0.272; pairwise comparisons: recent-control p = 0.01, historical-control p = 0.04; Fig. 5), but with no significant compositional differences between treatments (db-RDA, F = 1.533, p = 0.1; Fig. 5).

**Figure 5:**
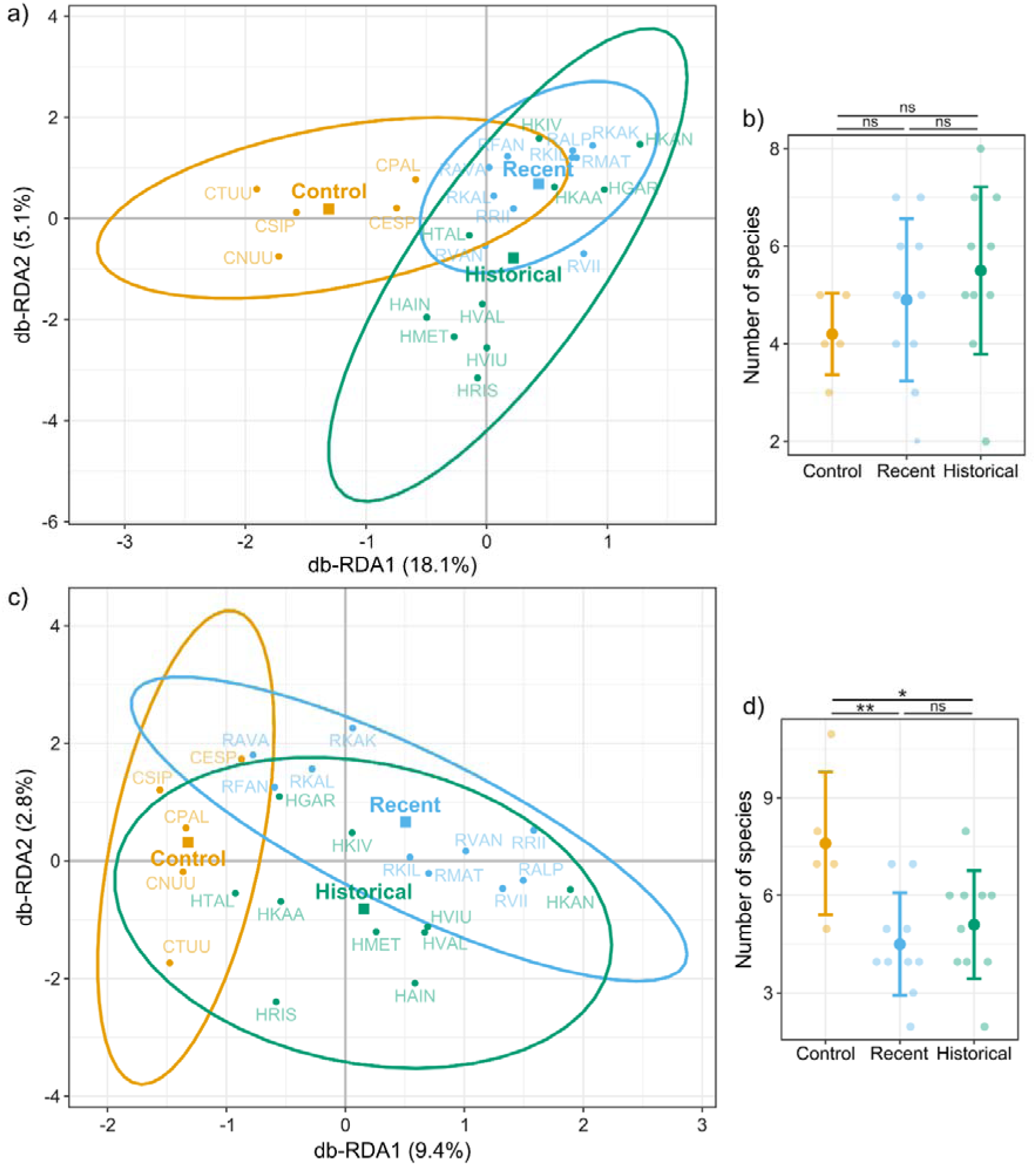
Distance-based Redundancy Analysis (db-RDA) showing the effect of treatment on a) long-winged, and c) short-winged species. Percentages on the axes indicate the proportion of variance explained by the constrained axes. Ellipses represent 95% confidence intervals. Number of species (± SD) per treatment for b) the long-winged, and d) the short-winged species.

### Species traits at the population level

We analysed responses of the 10 most abundant species to treatment, in terms of elytra length, mass corrected for size (both as indicators of animal quality and resource availability), and wing length corrected for size (proxy of dispersal capacity). Here, we highlight the most significant findings (Fig. 6, see all results in the Supplementary Materials, Figs. S2 to S4).

**Figure 6:**
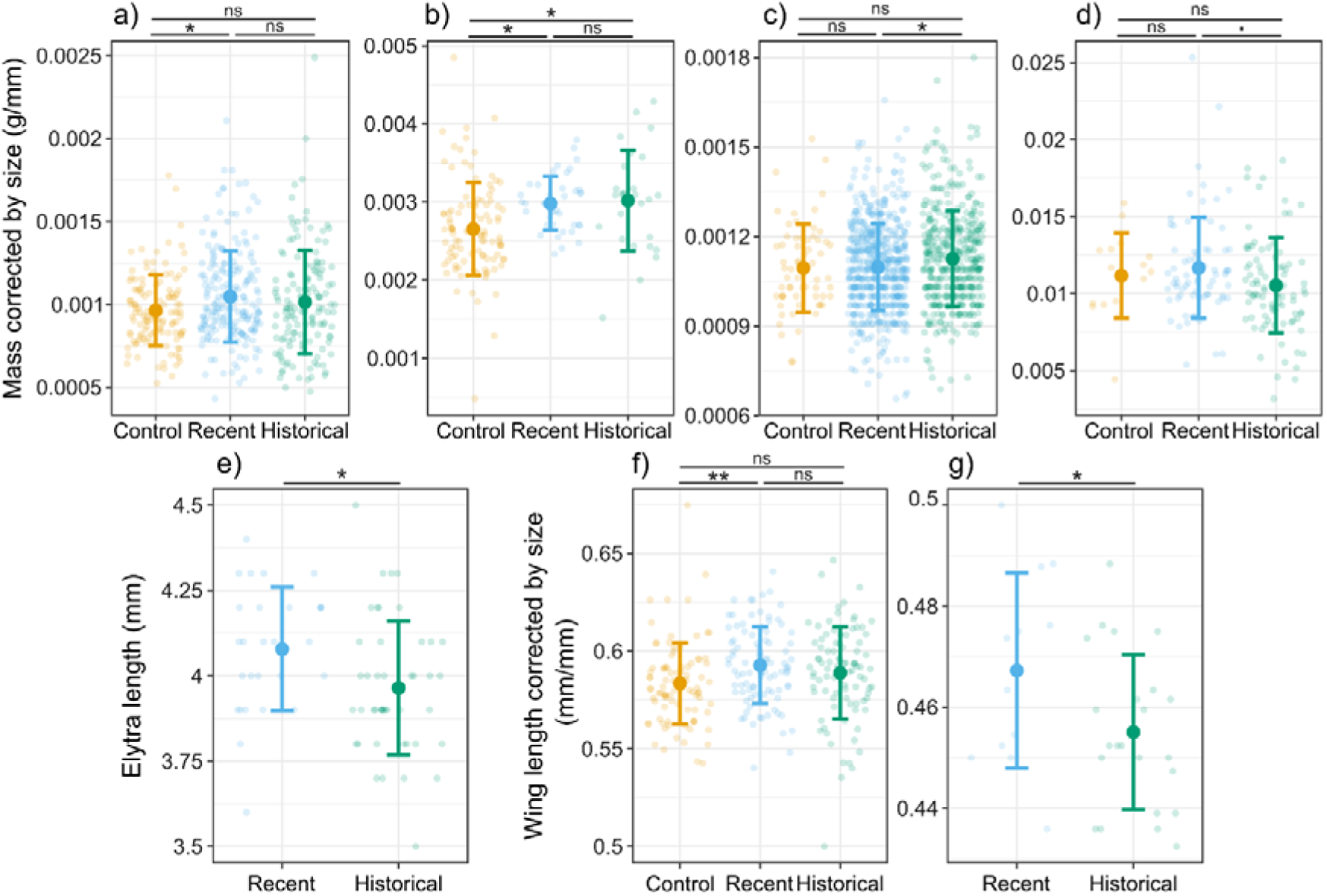
Mass corrected by size (± SD) per treatment for a) *Calathus micropterus*, b) *Pterostichus oblongopunctatus*, c) *Amara brunnea*, and d) *Carabus nemoralis*. e) Elytra length (± SD) per treatment for *Leistus ferrugineus.* Wing length corrected by size (± SD) per treatment for f) *Pterostichus niger*, and g) *Leistus ferrugineus*.

Both *C. micropterus* (Kruskal-Wallis, χ² = 6.5, p = 0.039; pairwise: recent-control, p = 0.026) and *P. oblogonpunctatus* individuals (LM, F = 7.847, p < 0.001; pairwise: historical-control, p = 0.008; recent-control, p = 0.005) were lighter in the control compared to the fragmented urban forests. *A. brunnea* was heavier (LM, F = 4.407, p = 0.012; pairwise: historical-recent, p = 0.011), while *C. nemoralis* was slightly lighter in the historically fragmented forest sites (LM, F = 2.695, p = 0.07; pairwise: historical-recent, p = 0.056). *L. ferrugineus* had longer elytra in the recently fragmented forests (LM, F = 6.21, p = 0.015).

*P. niger* had shorter wings (corrected for body size) in the control forests (LM, F = 4.424, p = 0.013; pairwise: recent-control, p = 0.009), while *L. fullegineus* had shorter wings in historically fragmented urban forests (LM, F = 4.187, p = 0.049).

## Discussion

We highlight the importance of considering both time and species traits when examining community responses to habitat fragmentation. In remnant urban forests of Helsinki, changes in carabid beetle communities occurred rapidly after fragmentation, yet we found no overall decline in richness. Instead, trajectories of different groups were strongly trait-dependent. Forest specialists exhibited a non-linear response, with richness declining sharply during the first three decades after fragmentation before recovering, while open-habitat species showed the opposite. Dispersal capacity proved decisive: species with limited dispersal declined in richness without compositional changes, whereas highly dispersive species maintained richness but underwent compositional shifts. These findings indicate that carabid beetles do not exhibit an extinction debt; responses are immediate and shaped by ecological specialisation and dispersal ability. At the population level, beetles showed no consistent pattern in size or dispersal traits, indicating that responses to fragmentation are highly species-specific and that, in the short to medium term, community-level changes are more sensitive indicators of fragmentation history than population-level traits.

Overall species richness did not differ between treatments. Even when comparing control, unfragmented forests to urban, fragmented forests, we observed no change in total species numbers. This contrasts with many studies reporting declines in diversity with habitat fragmentation, including ants (Carpintero & Reyes-López 2014), anurans (Herkert 1994; Ramalho *et al*. 2022), birds (Herkert 1994), and carabid beetles (Fujita *et al*. 2008; Koji *et al*. 2024; Niemelä 2001; Niemelä & Kotze 2009). In our study, the only factor positively influencing richness was temperature, highlighting that even within a northern city, small microclimatic variation can significantly affect diversity, consistent with the classic latitudinal gradient pattern in species richness (Pianka 1966). When considering cumulative richness at the landscape scale, however, a clear pattern emerges: older fragmented sites supported more species than both recent fragments and large control forests. This aligns with recent arguments that a mosaic of smaller, fragmented sites can maintain higher regional diversity than a single large forest (Fahrig 2017; Perrin *et al*. 2025; Riva *et al*. 2024), but the effect appears only over time. One plausible explanation is that urban fragmented forests attract species absent from large rural forests (which are generally quite species poor, see Niemelä et al. 2007), particularly open-habitat specialists, or species more adapted to a warmer environment due to the urban heat island effect. Consistently, community composition differed rapidly between fragmented and control sites. Historical sites were more variable than recent ones, suggesting that fragmentation initially filters communities, leaving only less vulnerable species, but over decades, community composition diversifies as pressures relax (McKinney 2006). Interestingly, the two large urban control forests (CPAL and CESP) were closer to fragmented sites in terms of community composition than the three rural controls. This indicates that even these large urban forests are beginning to exhibit community changes similar to fragmented sites, likely due to ongoing urban influence.

Focusing on key functional groups, such as habitat specialists and species with different dispersal capacities, reveals more nuanced patterns. These traits are known to strongly influence temporal responses to habitat changes (Chen *et al*. 2023). Forest specialists exhibited a non-linear dynamic in richness: a rapid decline during the first 30 years after fragmentation was followed by a gradual increase. Open-habitat species showed the opposite, initially increasing as forest edges and gaps created new habitat (see (Noreika & Kotze 2012), then declining. This temporary rise of open-habitat species mirrors the transient excess of rare species predicted by Hanski and Ovaskainen (2002), helping explain why overall species numbers remained relatively stable over time. A similar pattern emerged for body size, with fewer large species in recently fragmented sites. We observed no evidence of an extinction debt, as changes occurred quickly; however, there appears to be a “recovery debt” of roughly 30 years, with communities not returning to their initial state within our study period (Moreno-Mateos *et al*. 2017). While previous studies have documented the sensitivity of habitat specialists to fragmentation (Martinson & Raupp 2013), our results are the first to demonstrate a clearly non-linear dynamic in carabid beetles.

Dispersal capacity is a key trait influencing the temporal effects of habitat fragmentation (Finand *et al*. 2024; Niebuhr *et al*. 2015). Long-dispersal species (winged) exhibited changes in community composition following fragmentation, but their overall species richness remained stable. In contrast, short-dispersal species (wingless) experienced declines in richness without significant changes in composition. This pattern suggests that highly dispersive species can adapt by colonising suitable sites and abandoning less suitable ones, thereby altering community composition while maintaining species numbers. Low-dispersing species, however, are confined to their sites, leading to local extinctions of the most vulnerable species and no compensatory colonisation, which results in decreased richness but unchanged composition. Importantly, these changes occur rapidly, indicating the absence of an extinction debt for carabid beetles (Blowes *et al*. 2019; Ferraz *et al*. 2003). These findings complement our observations for habitat specialists, highlighting that trait-mediated responses strongly determine community dynamics after fragmentation (Keinath *et al*. 2017; Öckinger *et al*. 2010). Our study provides one of the first demonstrations of this pattern in short-lived, ground-dwelling insects. Few other studies have directly examined how dispersal capacity mediates fragmentation effects, though similar patterns have been reported in fungi, where low-dispersal species are more strongly affected by past fragmentation than highly dispersive species (Raimbault *et al*. 2024).

We observed no extinction debt for any species or functional group, confirming that short-lived carabid beetles respond rapidly to habitat changes. In contrast, most studies reporting extinction debt have focused on long-lived plants (Cousins 2009; Helm *et al*. 2005; Krauss *et al*. 2010; Lindborg & Eriksson 2004; Löffler *et al*. 2020; Vellend *et al*. 2006). Only a few studies have documented time-lagged responses in other taxa, such as birds (Almeida-Gomes *et al*. 2025), mammals (Ancillotto *et al*. 2025), pollinators (Cusser *et al*. 2015), or lichens (Hämäläinen & Fahrig 2024). Among insects, butterflies are one of the few groups showing a time lag (Löffler *et al*. 2020), but even this effect is not consistent across species, particularly specialists (Krauss *et al*. 2010). Our results highlight the importance of considering species traits when assessing temporal responses to fragmentation. Trait-based approaches reveal patterns that are obscured when looking only at overall species richness (Fontana *et al*. 2021).

In contrast to clear community-level patterns, population-level traits such as body size, mass, and dispersal ability showed no consistent response to fragmentation age. This contrasts with studies reporting body size reductions in urban or fragmented landscapes for carabids (Keinath *et al*. 2023; Papp *et al*. 2020) and other insects, including pollinators (Warzecha *et al*. 2016) and dung beetles (Bernardino *et al*. 2024; Franco *et al*. 2023). One explanation may be that urbanisation in Helsinki is less intense than in larger European capitals, resulting in weaker selective pressures. Alternatively, population-level traits may require finer-scale or longer-term monitoring to detect shifts.

Our study underscores the importance of looking beyond overall species richness and focusing on specific groups and traits. These trait-based responses have clear implications for conservation strategies. Forest fragmentation due to urbanisation produces immediate effects, without a detectable time lag, and fragmented forests do not fully recover to their original state, even after several decades. This highlights that decisions leading to fragmentation must be carefully considered, as their consequences are rapid, long-lasting, and often irreversible. Conservation efforts should prioritise vulnerable groups, such as habitat specialists and low-dispersal species, whose responses can be strong and sometimes non-linear. Understanding these trait-mediated effects is essential for predicting and mitigating the impacts of urban fragmentation on biodiversity. Extending this approach to other cities and taxa will be essential to test whether the rapid responses and “recovery debt” we observed are general features of short-lived organisms. Long-term monitoring that integrates community and population traits will also help refine conservation strategies in urban landscapes.

## Supporting information

Sup Mat

## Acknowledgements

We thank Sem de Waard for assistance with site selection and the City of Helsinki for authorising our sampling. We are grateful to URBARIA for funding Basile Finand’s salary. We also thank Sampsa Malmberg for species identification and Tessa Kajander, Heikki Setälä, Sara Turiel-Santos, Sini Ruohomäki and Dorottya Hajnal for their help in the field and laboratory. Finally, we acknowledge the use of AI for assistance with manuscript writing.

