## Supplementary material for "Habitat specialisation and dispersal capacity drive rapid carabid beetle responses to urban forest fragmentation": Sup Mat

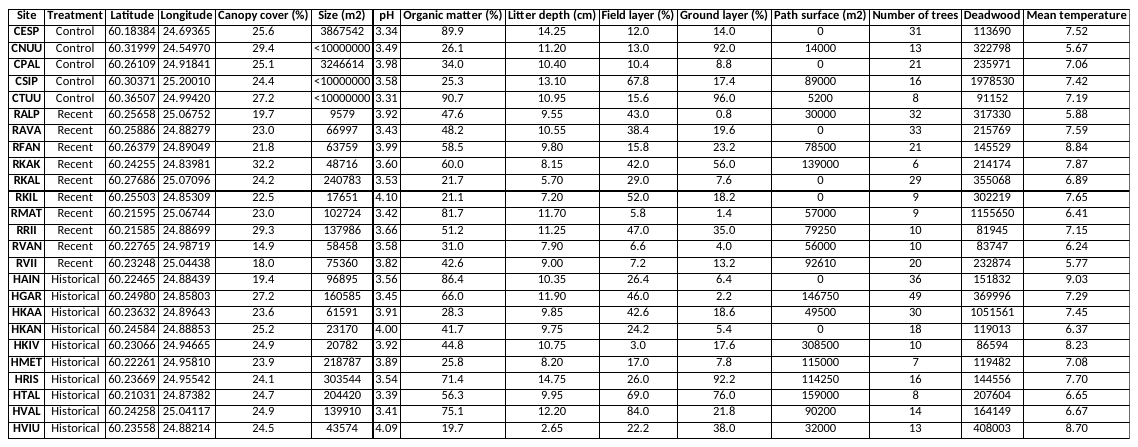

**Table S1:** Environmental characteristics measured for each site and coordinates.

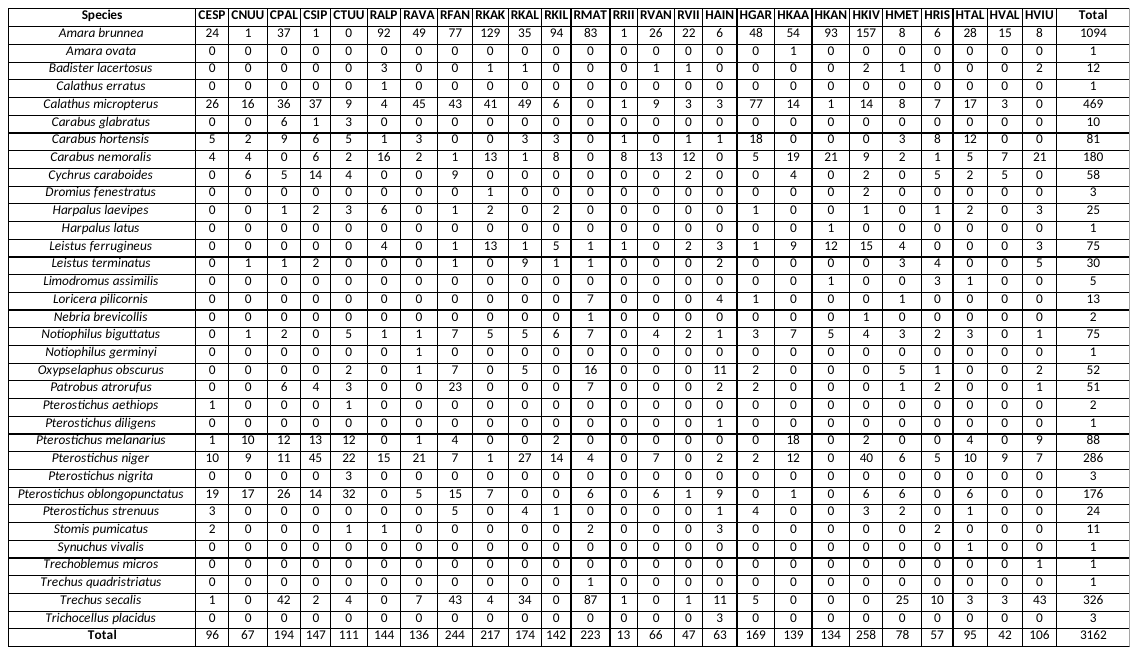

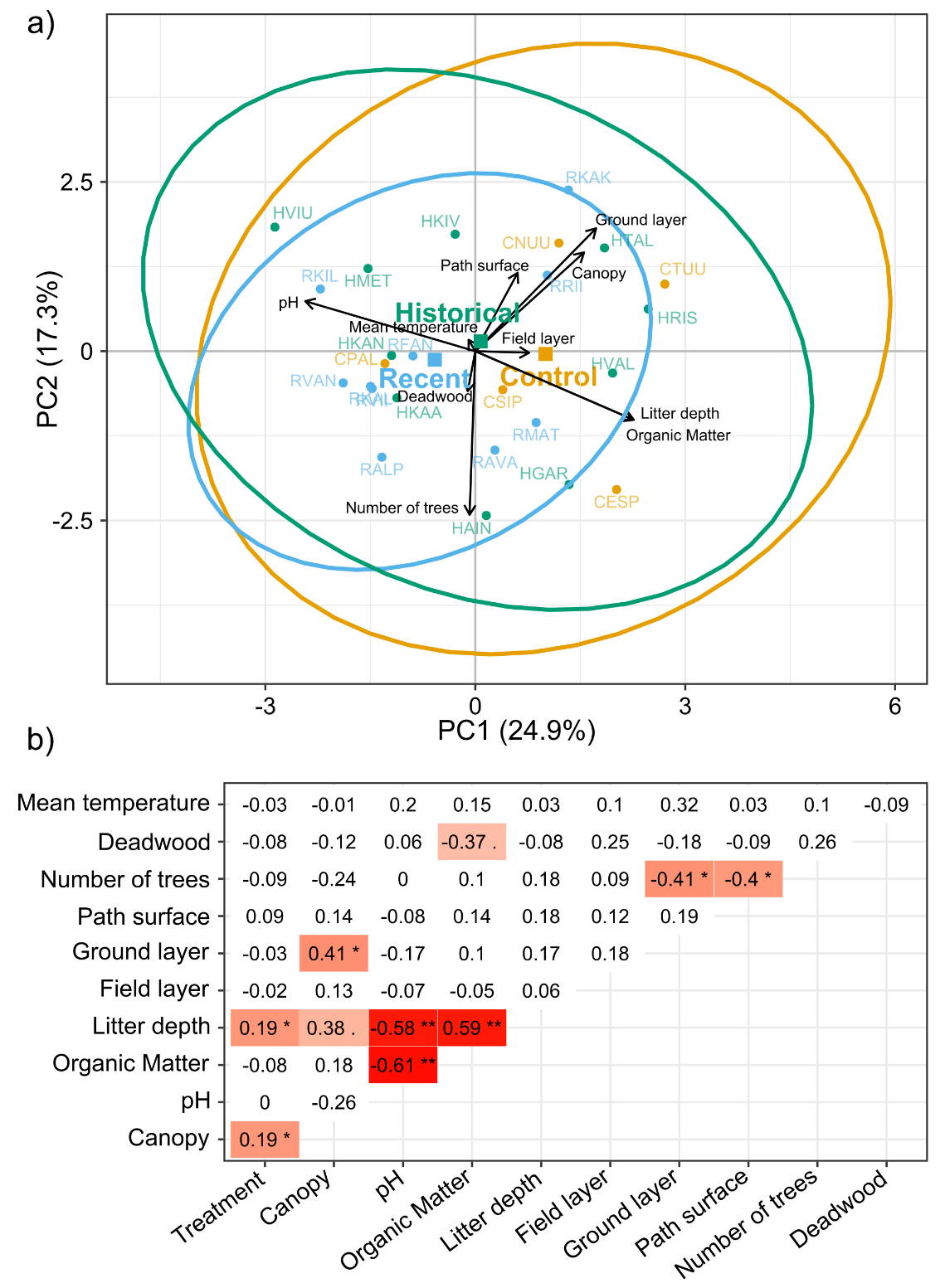
**Figure S1:** a) Principal Component Analysis (PCA) of environmental variables. Each point represents a site, and the ellipses indicate 95% confidence intervals for the three treatments (Historical, Recent, Control). Arrows represent the direction and relative strength of each environmental variable. The percentage on each axis shows the proportion of variance explained by that principal component. b) Correlation matrix of environmental variables. Each value represents the correlation coefficient, with the sign indicating the direction of the relationship. Colour intensity and stars indicate the significance of the p-value: a dot (.) indicates p < 0.1, * indicates p < 0.05, and ** indicates p < 0.01. Darker colours correspond to stronger correlations and higher significance.

**Table S2:** List of species per site. The values represent the number of individuals per species collected from this study.

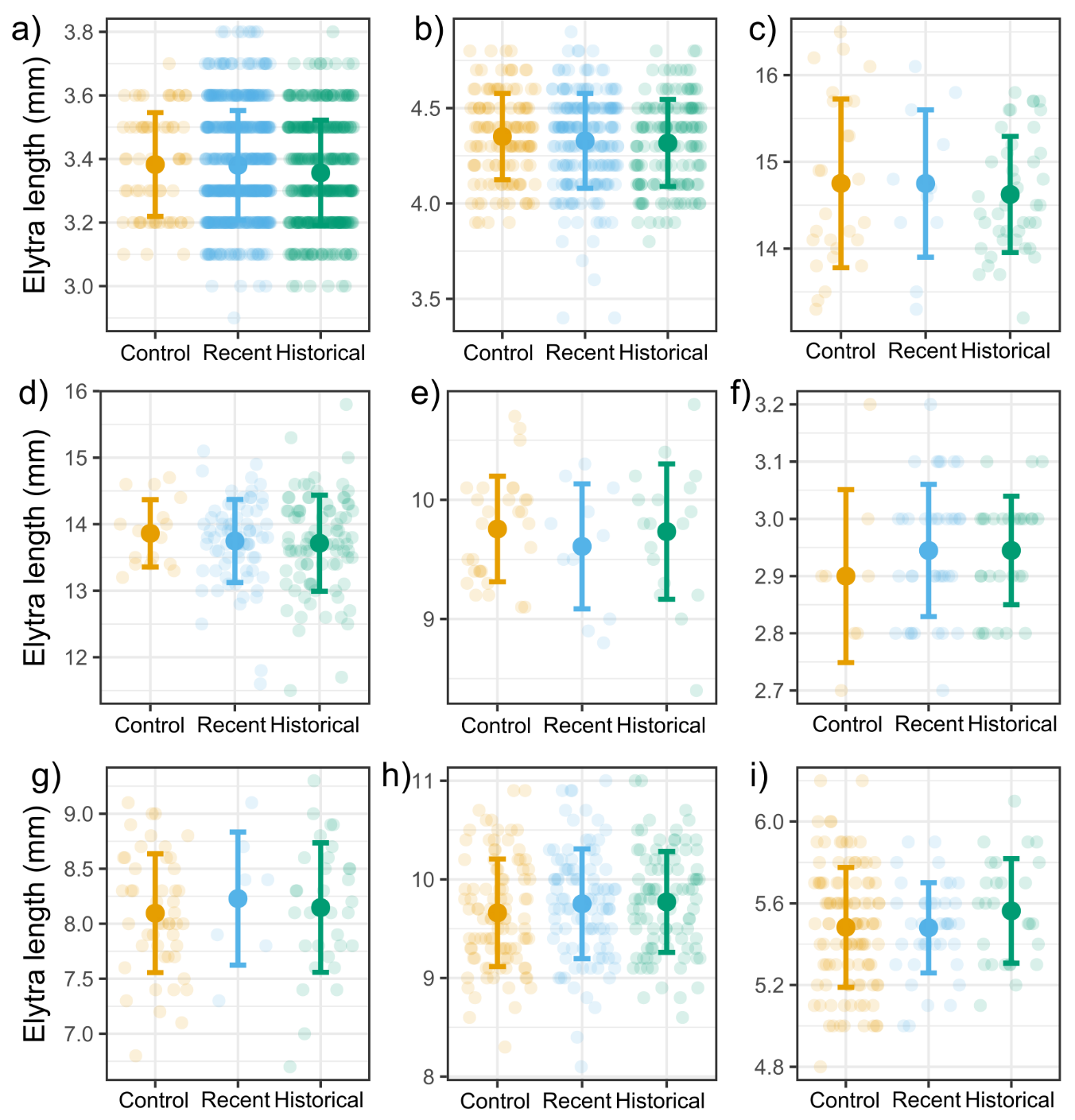

**Figure S2:** Elytra length (± SD) per treatment for a) *Amara brunnea,* b) *Calathus micropterus,* c) *Carabus hortensis*, d) *Carabus nemoralis,* e) *Cychrus caraboides*, f) *Notiophilus biguttatus*, g) *Pterostichus melanarius,* h) *Pterostichus niger*, and i) *Pterostichus oblongopunctatus.*

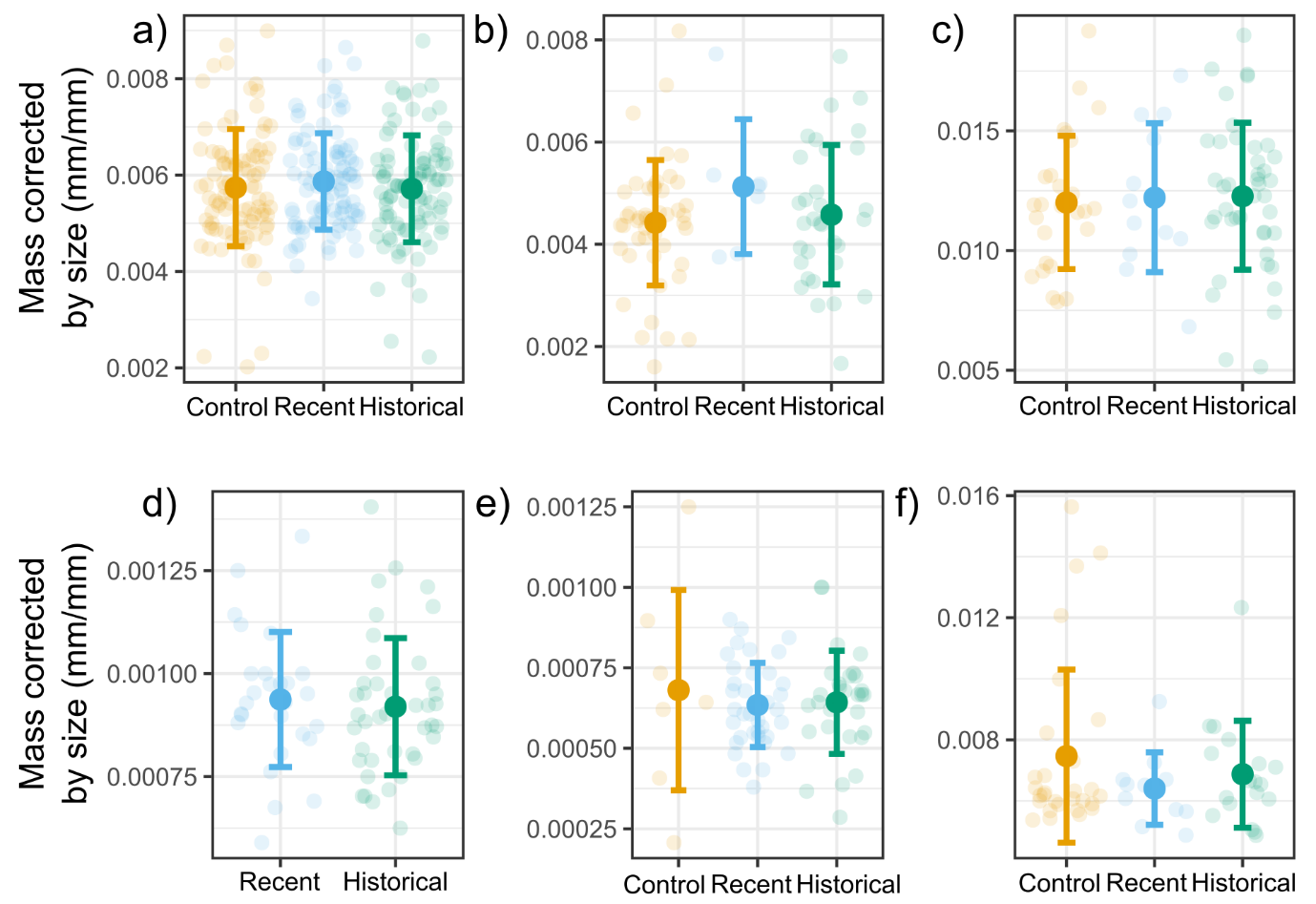
**Figure S3:** Mass corrected by size (± SD) per treatment for a) *Pterostichus niger,* b) *Pterostichus melanarius,* c) *Carabus hortensis, d) Leistus ferrugineus,* e) *Notiophilus biguttatus,* and f) *Cychrus caraboides.*

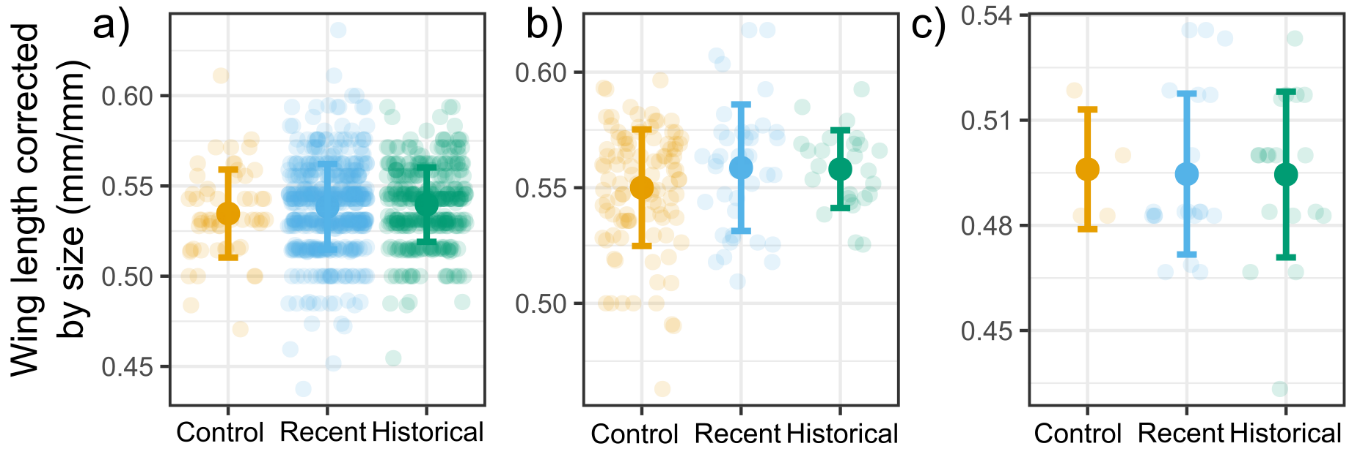

**Figure S4:** Wing length corrected by size (± SD) per treatment for a) *Amara brunnea,* b) *Pterostichus oblongopunctatus,* and c) *Notiophilus biguttatus.*
